# Behavioral evidence for visually dominant audiovisual motion integration

**DOI:** 10.1101/2025.10.07.680959

**Authors:** Adam J. Tiesman, Kalina Stoyanova, Ramnarayan Ramachandran, Mark T. Wallace

## Abstract

Accurate motion perception is crucial for navigating complex environments, where sensory information is often derived from multiple modalities. While multisensory cues improve performance in tasks involving stationary stimuli, the integration of dynamic motion cues is less understood. Here, we examined the nature of auditory and visual motion integration and determined whether attentional instructions modulate integration. Sixty participants performed direction discrimination tasks under auditory, visual, and audiovisual conditions, with cueing instructions to attend to one or both modalities. Results showed that visual motion direction estimates were more reliable than auditory ones, and audiovisual estimates were superior to either modality alone. Additionally, visual motion estimates were more stable that auditory motion estimates during direction conflict trials. Reliability-weighted models of integration closely predicted audiovisual performance, demonstrating optimal integration. Sensory dominance (visual vs. auditory) influenced sensitivity to audiovisual motion, with visual dominant participants performing better overall. Our results demonstrate that the perception of audiovisual motion is dominated by vision and is shaped by top-down attention.

## 1 Introduction

Perceiving and responding to motion are critical for survival, shaping behaviors such as evasion, foraging, and navigation. To support these kinds of actions, the brain contains specialized mechanisms for processing motion. In the primate visual system, motion-selective neurons have been extensively characterized in cortical areas including the middle temporal (MT) and medial superior temporal (MST) areas, where responses are tuned to direction of visual motion and where cortical damage produces deficits in motion perception [Andersen, 1997b, Maunsell and Van Essen, 1983]. In contrast, for the auditory system, there is surprisingly little evidence for dedicated spatially selective motion areas [Poirier et al., 2005, Chaplin et al., 2018b]. Although humans can perceive auditory motion, it is typically thought to be derived from dynamic interaural time and level differences rather than directly mapped in spatial coordinates [Carlile and Leung, 2016].

Importantly, objects moving within out everyday environments are rarely unisensory. Many natural events generate concomitant auditory and visual motion cues. It is natural to ask how the brain actively integrates these cross-modal signals and how that integration impacts behavior to those events. Multisensory integration has been well documented in other domains, including speech perception, object recognition, and spatial localization [Wallace et al., 2004, Stevenson et al., 2014]. Although there has been some work investigating whether auditory motion cues influence visual motion cues [Soto-Faraco et al., 2003, Meyer and Wuerger, 2001, McCourt and Leone, 2016], there is little to no conclusive evidence that audiovisual motion information confers a perceptual benefit beyond what is available through audition or vision alone.

Prior work involving static cues has established that when multiple sensory cues are available, they are typically combined in a statistically optimal manner. Using reliability-weighted models of integration such as maximum likelihood estimation (MLE), behavioral and neural evidence support that the nervous system weights each sensory modality according to its relative reliability to minimize variance in the multisensory estimate [Ernst and Banks, 2002, Alais and Burr, 2004b, Morgan et al., 2008, Battaglia et al., 2003]. This variance minimization allows the nervous system to form the most precise estimate of a multisensory object, provided noisy sensory information. Some work has found neural correlates of this model for the integration of visual and vestibular motion cues [Angelaki et al., 2009, Fetsch et al., 2011]; however, little work has investigated whether this reliability-weighted model holds true for auditory and visual motion cue integration.

Furthermore, multisensory processing is not purely bottom-up; it can be modulated by top-down mechanisms such as expectations, reward, learning, and attention [Kayser and Kayser, 2020, Duyar et al., 2022, Talsma et al., 2010, Donohue et al., 2011, Busse et al., 2005, Blurton et al., 2015]. Cueing participants to attend to a specific sensory modality can alter both the way sensory information is weighted (via reliability-weighted integration models) and how it is combined across modalities to give rise to an audiovisual estimate [Rohe and Noppeney, 2018]. The role of attention in multisensory motion perception remains particularly underexplored, specifically how attention alters audiovisual estimates of motion.

In the present study, we asked how audiovisual motion stimuli are combined using a motion discrimination paradigm. We also examined whether the allocation of attention (via cueing) modulates this integration. Using psychometric curve fitting and reliability-weighted modeling, we compared motion direction sensitivity across unisensory and multisensory conditions (see Figure 1 for an overview). Intuition led us to surmise that participants would integrate auditory and visual motion cues in a reliability-weighted fashion, and that attention to a given modality would modulate discrimination performance.

**Figure 1.**
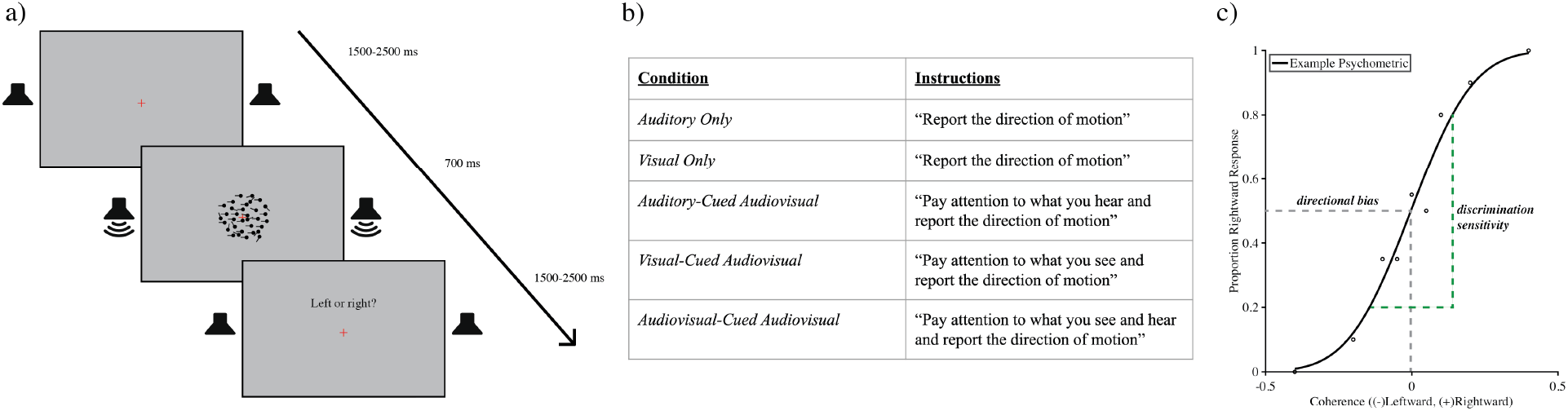
Experimental design. (a) For all conditions, participants were instructed to fixate on a fixation cross in the middle of the screen, followed by a 700 ms A, V, or AV stimulus presentation, and ending with a jittered response window. (b) Task conditions all participants were tested on, with instructions being the only difference in the multisensory conditions. (c) Discrimination sensitivity (slope of the psychometric curve) was the main metric used to assess performance across our task conditions.

## 2 Results

### 2.1 Unisensory Performance

Discrimination sensitivities (slope of the psychometric function) for auditory only and visual only conditions are shown in Figure 2a. At a group level, the visual only sensitivities were significantly higher than the auditory only sensitivities (*p* = 0.0254). These sensitivities were compared within individuals to assess the relative performance of auditory only to visual only estimates of motion (Figure 2b). Individual variability in unisensory discrimination sensitivity was high, with some participants showing greater auditory sensitivity (i.e., data points below the diagonal line: auditory dominant) and others showing greater visual sensitivity (i.e., data points above the diagonal line: visual dominant). Exemplar psychometric curves of two participants showcasing either a greater auditory only or visual only sensitivity are shown (Figure 2c). For our task, there were more visual dominant participants (n=30) compared to auditory dominant participants (n=18). This all suggests that visual motion estimates are generally more reliable for this task.

**Figure 2.**
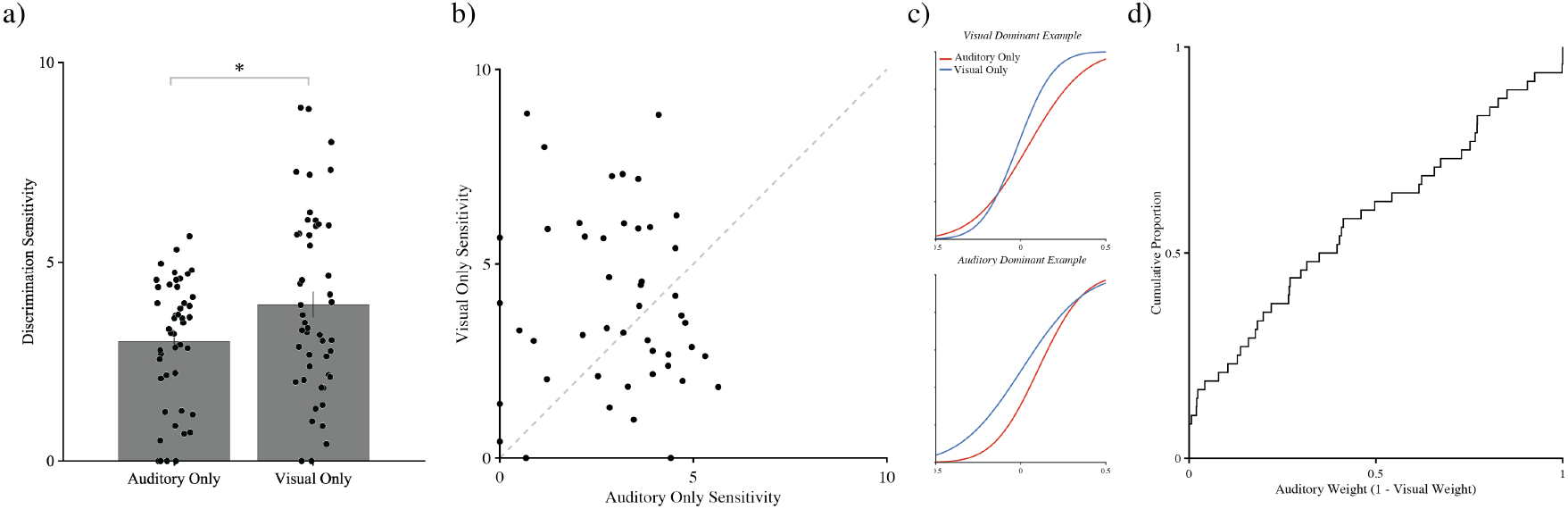
Unisensory auditory and visual motion judgments. (a) Discrimination sensitivities for both the auditory and visual only conditions are shown. (b) Relative unisensory discrimination sensitivity is shown with (c) psychometric curves showing example visual dominant (top) and auditory dominant (bottom) participants. (d) An empirical cumulative distribution function showing the group distribution of predicted sensory weights using maximum likelihood estimation is shown.

We then calculated sensory weights using maximum likelihood estimation (MLE). Under the assumptions of MLE, these sensory weights reflect the relative reliability of each modality in forming a multisensory estimate. A higher auditory weight, for example, indicates that a participant relies more on the auditory cue than the visual cue when both are presented together [Ernst and Banks, 2002]. Similar to the variability observed in unisensory sensitivities, the predicted sensory weights varied substantially across participants. An empirical cumulative distribution function of all participants and their sensory weights is shown in Figure 2d. For these sensory weights, participants who had an auditory weight greater than 0.5 would be predicted to have their audiovisual percept of motion “dominated” by audition. Likewise, participants with an auditory weight of less than 0.5 would be predicted to have their audiovisual percept of motion “dominated” by vision. This measure was then used to split the group into two cohorts: auditory dominant (AD) and visual dominant (VD). These cohorts were used to further evaluate differences in sensory dominance and how this impacts audiovisual motion performance. This heterogeneity suggests that the perception of audiovisual motion is highly individualized; audiovisual motion for many participants is disproportionately captured by one sensory modality. In uncued audiovisual conditions, we could not control for this individual sensory dominance, meaning that participants’ judgments on motion would likely be driven by one sensory modality, and the results would seldom show any multisensory enhancement [Alais and Burr, 2004a, Ernst and Bulthoff, 2004]. To address this we implemented cued multisensory conditions in which attention was explicitly directed towards audition, vision, or both. This approach allowed us to control decision strategy, thereby directing participants to evaluate audiovisual motion through the use of auditory and visual motion cues.

### 2.2 Impact of Cueing on Performance on Congruent Multisensory Trials

Judgments about audiovisual motion on cued trials were then compared to unisensory performance for both AD and VD participants. Mean discrimination sensitivity for each condition was plotted for both AD and VD participants. First, we looked at auditory conditions (i.e., conditions where participants were instructed to report the auditory motion). For both AD and VD participants, when comparing auditory only to auditory cued AV condition, there was a significant improvement in discrimination sensitivity (Wilcoxon signed-rank: *p* = 0.0234, *p* = 0.0014). However, when subsequently comparing auditory cued AV to audiovisual cued AV conditions, only VD participants showed an increase in discrimination sensitivity (Wilcoxon signed-rank: *p* = 0.0027) (Figure 3a). It is important to note that the variability in auditory cued AV and audiovisual cued AV sensitivities was large for both groups (*σ* = 2.67, *σ* = 3.55). Therefore, some AD participants may have exhibited multisensory enhancement, and some VD participants may have not shown any enhancement. However, at a group level, when comparing auditory only conditions to cued multisensory conditions, only VD participants seem to show consistent improvement in discrimination sensitivities.

**Figure 3.**
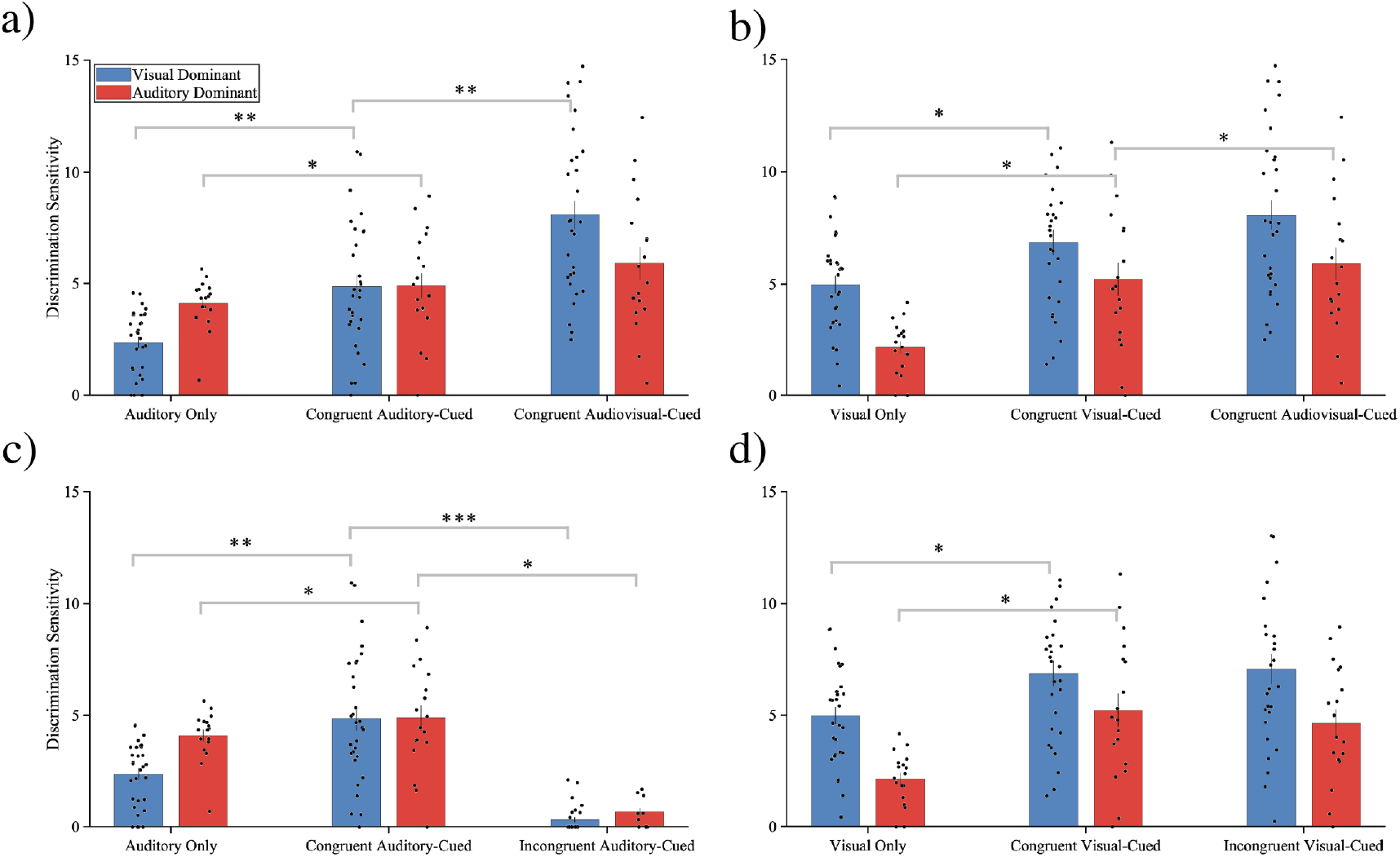
Congruent and incongruent multisensory judgments of motion. (a) Discrimination sensitivities for auditory conditions (i.e., auditory only, auditory cued AV, and audiovisual cued AV) are shown, separated by AD and VD participants. (b) Discrimination sensitivities for visual conditions (i.e., visual only, visual cued AV, and audiovisual cued AV) are shown, separated by AD and VD participants. (c) Auditory motion judgments for auditory only, congruent auditory cued, and incongruent auditory cued conditions are shown, separated by AD and VD participants. (d) Visual motion judgments for visual only, congruent visual cued, and incongruent visual cued conditions are shown, separated by AD and VD participants.

We then examined visual conditions (i.e., conditions where participants were instructed to report the direction of the visual motion). It is important to note that for the audiovisual cued AV condition, participants were instructed to attend to both the auditory and visual cues, hence, this condition is shown in both Figure 3a and 3b. AD participants had significantly higher visual cued AV and audiovisual cued AV sensitivities compared to their visual only sensitivities (Wilcoxon signed-rank: *p* = 0.0504, *p* = 0.0078). Similarly, for the VD participants, visual cued AV and audiovisual cued AV sensitivities were significantly higher than visual only sensitivities (Wilcoxon signed-rank: *p* = 0.0186, *p* = 0.0081). For the visual cued AV condition, VD participants had significantly higher discrimination sensitivities compared to AD participants (Wilcoxon signed-rank: *p* = 0.0481). However, for both the auditory cued and audiovisual cued AV condition, the AD and VD group sensitivities were not significantly different (Wilcoxon signed-rank: *p* = 0.064, *p* = 0.411). VD participants seem to both be exhibiting more multisensory enhancement as well as showing higher sensitivity across most multisensory conditions compared to AD participants. VD participants on average have better estimates of multisensory motion direction, especially when cued towards vision. These results reveal that sensory weighting predicts performance for cued audiovisual motion judgments, as performance of VD participants for visual cued and audiovisual cued AV conditions showed the best audiovisual task performance.

### 2.3 Performance Across Incongruent Trials

For all audiovisual motion judgments, 20% of the trials were incongruent, meaning the auditory and visual stimuli moved in opposite directions. Performance on these trials was compared against performance on both unisensory and cued congruent AV trials. For auditory conditions (i.e., conditions where participants were instructed to report the auditory motion), both AD and VD participants showed significantly lower discrimination sensitivities for incongruent auditory cued AV trials compared to both auditory only (Wilcoxon signed-rank: *p* = 0.00781, *p <* 0.001) and congruent auditory cued AV trials (Wilcoxon signed-rank: *p* = 0.0156, *p <* 0.001) (Figure 3c). For incongruent auditory cued AV trials, both AD and VD participants showed similarly poor performance (Wilcoxon signed-rank: *p* = 0.449) (Figure 3c). For incongruent auditory cued AV trials, both groups suffer in discrimination sensitivity, suggesting that conflicting visual motion information hinders auditory motion direction discrimination for our task.

For visual conditions (i.e., conditions where participants were instructed to attend to the visual motion), we again compared incongruent AV to unisensory and congruent AV performance. AD and VD participants showed significantly higher discrimination sensitivities for incongruent visual cued AV trials compared to visual only trials (Wilcoxon signed-rank: *p* = 0.039, *p* = 0.0055) but not compared to congruent visual cued AV trials (Wilcoxon signed-rank: *p* = 0.461, *p* = 0.446) (Figure 3d). VD participants had trending higher discrimination sensitivities in incongruent visual cued AV trials compared to AD participants (Wilcoxon signed-rank: *p* = 0.0784) (Figure 3d). It is important to note that there was an overall low number of incongruent trials, which limits the confidence with which strong claims about the impact of incongruence on auditory and visual motion judgments. We do, however, want to highlight that incongruent visual cued AV judgments seem to be less affected by conflict compared to incongruent auditory cued AV judgments. This is seen for both AD and VD participants, and the differences between AD and VD participants may point to differences in cross-modal conflict processing. Possibly, VD participants are better able to segregate conflicting crossmodal signals to facilitate accurate judgments of visual motion direction.

### 2.4 Reliability-Weighted Integration of Motion

Using the unisensory performance, the predicted AV sensitivities were derived using MLE, a reliability-weighted integration model (Equation 4). The three cued congruent AV conditions were then compared to these MLE predictions, split by AD and VD participants. The MLE predicted AV discrimination sensitivities for AD and VD participants were not significantly different (Wilcoxon signed-rank: *p* = 0.222) (Figure 4a). First, for AD participants, discrimination sensitivities in all multisensory conditions (auditory cued, visual cued, and audiovisual cued) did not significantly differ from MLE predictions (Wilcoxon signed-rank: *p* = 0.753, *p* = 0.641, *p* = 0.547) (Figure 4a). Similarly, for VD participants, discrimination sensitivities in all multisensory conditions did not significantly differ from MLE predictions (Wilcoxon signed-rank: *p* = 0.355, *p* = 0.1011, *p* = 0.0619) (Figure 4a). However, for VD participants, it is important to note that the multisensory cued sensitivities were trending towards being higher than the estimated MLE sensitivities. This led us to explore other factors that may explain the observed, albeit trending, deviation from the reliability-weighted model prediction. And as such, these results suggest that MLE accounts well for AD participants’ judgments across cued AV conditions, but does not fully capture the patterns observed in VD participants’ cued AV judgments.

**Figure 4.**
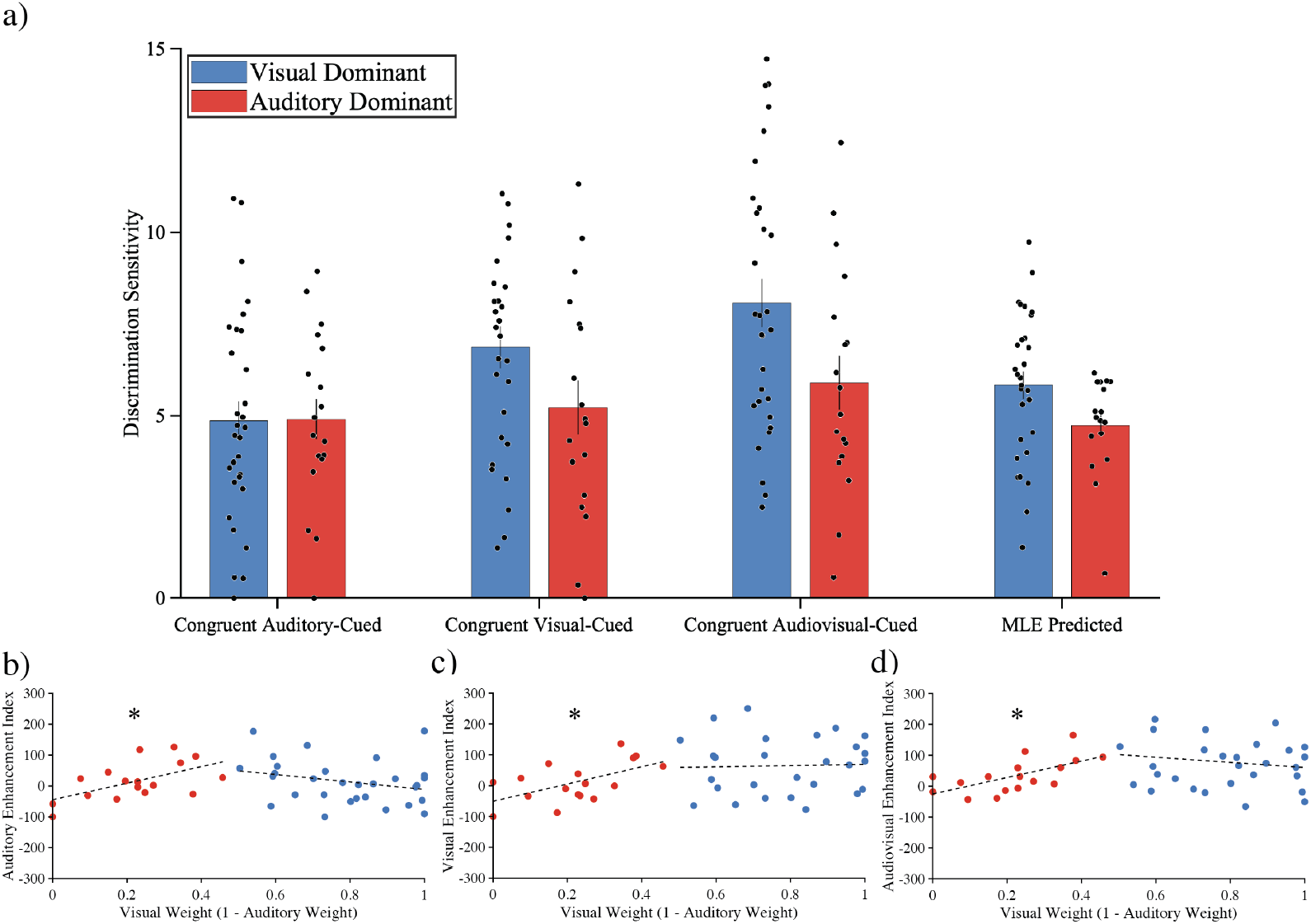
Comparing audiovisual performance against maximum likelihood estimation (MLE). (a) Auditory cued, visual cued, and audiovisual cued performance is compared against MLE predicted performance, split by AD and VD participants. (b) Separate linear regressions of sensory weights predicted by MLE and the auditory cued enhancement index measure for both AD and VD participants are shown. (c) Separate linear regressions of sensory weights predicted by MLE and the visual cued enhancement index measure for both AD and VD participants are shown. (d) Separate linear regressions of sensory weights predicted by MLE and the audiovisual cued enhancement index measure for both AD and VD participants are shown.

Equal sensory weighting should optimize sensory conditions for integration [Ernst and Bulthoff, 2004]. In other words, as two cues approach equal reliability (i.e., sensory weights for both approach 0.5), the enhancement in multisensory conditions should be the greatest. This relationship should hold true in conditions where participants are engaging in reliability-weighted integration (MLE). To test this, we ran two linear regressions, one for AD and one for VD participants, for each cued AV condition using each participant’s enhancement index and sensory weights. This enhancement index (EI) was calculated using Equation 5, where each multisensory condition (auditory cued, visual cued, audiovisual cued) was compared against a given participant’s best unisensory condition to give rise to three enhancement indices: auditory EI, visual EI, and audiovisual EI. We ran a total of six regressions across all multisensory conditions for both AD and VD participants (Figure 4b-d). Whereas AD participants had a significant correlation across cued AV conditions (*R*^2^ = 0.326, *p* = 0.017; *R*^2^ = 0.325, *p* = 0.017; *R*^2^ = 0.395, *p* = 0.007), VD participants failed to show any significant correlation in any cued AV condition (*p* = 0.148, *p* = 0.860, *p* = 0.454). Although none of the cued AV conditions significantly deviated from the reliability-weighted model’s predictions, for VD participants, performance for our task may not be entirely explained by these reliability-weighted integration models.

## 3 Discussion

In this study, we examined how auditory and visual cues are integrated during motion direction discrimination. Our results show that visual motion estimates were generally more reliable than auditory motion estimates, and sensory weighting led to differences in cued AV performance. We observed substantial between-subjects variability in performance, both within and across conditions (Figure 2). The heterogeneity in sensory weighting may reflect differences in prior sensory experience, auditory or visual motion expertise, differing cognitive strategies, as well as underlying differences in neural mechanisms. Future work could investigate the plasticity of these differences through training and assess whether such dominance remains stable over time within individuals.

The observed advantage for visual motion direction discrimination is consistent with the modality appropriateness hypothesis, which posits that the modality most suited to the task domain will dominate perception [Welch and Warren, 1980, Soto-Faraco et al., 2004]. Because direction discrimination relies largely on spatial information compared to temporal information, vision provides a more precise representation of motion than audition. Vision is supported by high spatial acuity and retinotopic mapping, whereas audition must infer spatial trajectories from interaural timing, level difference, and spectral cues [Carlile and Leung, 2016, Krumbholz et al., 2005]. However, different features of motion rely more on audition. A study by Freeman et al. found that human participants were most sensitive to auditory duration, a temporal feature of motion, compared to displacement or speed. Although participants were able to discriminate the speed of auditory motion, performance was worse when discriminating the duration of auditory motion [Freeman et al., 2014]. Future research could complement these findings by investigating parametric manipulations of both auditory-favored features and visual-favored features of motion to better understand how each sensory modality contributes to the perception of audiovisual motion.

It is important to consider that naturalistic audiovisual motion is rarely just in the azimuthal (i.e., horizontal) plane. The current study investigated the plane of space where the auditory system can most effectively utilize interaural time and level differences to infer motion trajectories. However, vertical motion trajectories in the auditory system cannot utilize binaural cues, but instead rely on monaural spectral cues, making them less reliable [Carlile and Leung, 2016, Battal et al., 2019, Britten et al., 1992, Snowden and Freeman, 2004]. Additionally, studies have found differences in multisensory facilitation between looming and receding audiovisual motion, adding to the complexity of multisensory motion perception [Cappe et al., 2009, 2012]. Therefore, investigating more ecologically valid audiovisual motion that combines horizontal, vertical, and looming/receding motion is necessary to elucidate how the brain dynamically represents and integrates motion information.

Cued AV conditions significantly enhanced discrimination sensitivity compared to unisensory conditions. This effect likely reflects both bottom-up sensory processing as well as top-down attentional control. Attended cues have more reliable sensory estimates compared to unattended cues in the same modality [Rohe and Noppeney, 2018]. Therefore, it is important to consider that across our multisensory conditions, the relative contribution of auditory and visual motion information differed, even though the stimulus statistics remained identical. As such, the predicted sensory weighting is for a multisensory condition that is assumed to be in the absence of top-down mechanisms such as attention and purely driven by bottom-up sensory integration (Figure 2). Our current task does not allow us to calculate empirical sensory weights and compare them to the model’s predicted sensory weights. Future work could investigate this direct comparison of empirical vs predicted weights using our audiovisual motion discrimination paradigm to more directly interrogate how top-down mechanisms like attention shape sensory estimates.

Notably, our cued AV conditions showed different integration profiles depending on the sensory dominance (AD and VD participants). For AD participants, all cued AV conditions were well predicted by reliability-weighted integration, with approximately 35% of audiovisual performance explained by the model’s sensory weighting. In contrast, for VD participants, these cued AV conditions were not fully explained by the same model. Although their performance did not significantly differ from the model’s predictions, their sensory weighting could not account for their audiovisual performance in our task (Figure 4b-d). One consideration for this is the contribution of attention to the model. Reliability-weighted models of integration do not assume any top-down contributions to the precision of the sensory estimates. As such, one simple extension to this model is having a third distribution that accounts for attention, represented by cueing, in our task [Ernst and Bulthoff, 2004]. If this were to be considered, possibly all conditions for our task would be optimal or suboptimal integration, consistent with previous studies attempting to characterize behavioral performance of motion perception [Alais and Burr, 2004a]. To our knowledge, our study is the first to find audiovisual motion perception that largely aligns with reliability-weighted integration models. Future work should corroborate these findings by investigating whether these behavioral models hold under different attentional loads and further develop extensions to these integration models.

An additional consideration is that effective integration depends not only on spatial and temporal coincidence but also on semantic congruence between cues. While the visual stimulus (dot clouds) implied motion direction without global displacement, the auditory stimulus contained explicit displacement cues. This asymmetry may have disrupted perceptual binding, or the unification of sensory cues into a single perceptual event. Although it has been posited that sensory binding is primarily driven by temporal coincidence, the nervous system is increasingly sensitive to stimulus semantics as stimulus complexity increases [Bizley et al., 2016]. Doppler effects, spectral cues, and timing differences across the two speakers were not used in the generation of our auditory stimuli. These disparities across the sensory modalities may create suboptimal sensory integration conditions. A leading theory of auditory motion perception is the “snapshot” model, where auditory motion information is inferred through a series of displacement cues [Carlile and Leung, 2016, Krumbholz et al., 2005]. If there is no visual displacement information available, there might not be perceptual binding of our stimuli, as the dominant sensory cues are crossmodally disparate (velocity for vision and displacement for audition). Instead of perceptual binding in multisensory motion conditions, participants may have resorted to switching their attention between vision and audition. Our results reveal significant multisensory enhancement despite the considerations described above (Figure 3). Future work could move towards simulating auditory and visual motion as naturally as possible to understand which features allow for the “natural” perceptual binding of a moving audiovisual object.

Incongruent trial performance offers preliminary insight into this crossmodal issue. Cueing participants to a specific sensory modality while the cues are conflicting allow us to explore if participants can appropriately segregate motion signals. For both AD and VD participants, incongruent auditory cued AV trials resulted in the lowest performance out of all conditions tested, in agreement with other audiovisual motion work [Soto-Faraco et al., 2004]. This suggests both that optimal direction discrimination performance requires visual information and that incongruent auditory information is more easily ignored when accompanied by visual motion. This is further supported by the fact that incongruent visual cued AV trials did not perturb discrimination sensitivity. Although we cannot directly assess this from these data, these observations raise the question of whether participants could discriminate between congruent and incongruent motion. If congruent and incongruent motion cues are discriminable, it would support the hypothesis that perceptual binding is occurring appropriately. Testing this explicitly is a key next step to disentangle whether multisensory enhancement is driven by perceptual binding (i.e., integration) or from performance improvements attributable solely to attentional cueing or cue dominance.

The behavioral patterns observed in this study suggest contributions from multiple neural substrates involved in motion processing, multisensory integration, and attentional control. Visual motion cues likely engage areas such as the human middle temporal complex (hMT+) [Rezk et al., 2020, Kayser et al., 2007, 2017], analogous to the non-human primate medial superior temporal (MST) and middle temporal (MT) areas, which are known for their role in responding to visual motion trajectories (i.e., direction and velocity) [Maunsell and Van Essen, 1983]. Area hMT+, or analogous areas MT/MST, may be recruited for the perception of multisensory motion in addition to its role in vision motion alone [Hagen et al., 2002, Chaplin et al., 2018b,a]. Multisensory integration of auditory and visual motion cues may be supported by convergence zones including the posterior parietal cortex and the superior temporal sulcus, regions previously implicated in combining spatial and temporal information across modalities [Molholm et al., 2006, Hocking and Price, 2008, Andersen, 1997a, Stevenson and James, 2009, Werner and Noppeney, 2010]. Early sensory areas such as primary auditory and visual cortices also may represent crossmodal information [Kayser et al., 2007, Kamitani and Tong, 2006, Alink et al., 2012]. Specifically, these regions may increase their connectivity when viewing congruent auditory and visual motion cues [Lewis and Noppeney, 2010, Alink et al., 2008]. In addition, the attentional effects observed, particularly the enhancement of performance in cued conditions, point to the involvement of top-down control networks, such as the dorsal attention network, which mediate voluntary attentional allocation, specifically for spatial tasks [Majerus et al., 2018, Rajan et al., 2021, Ptak and Schnider, 2010, Bremmer et al., 2001]. To test these hypotheses more directly, future studies should incorporate functional neuroimaging (e.g., fMRI) or electrophysiological (e.g., EEG, MEG) methods to measure neural responses specific to both the sensory encoding of dynamic cues as well as the executive control of decision making and modality-driven attention.

In summary, our findings demonstrate that visual motion estimates are generally more reliable than auditory motion estimates, yet audiovisual integration still enhances direction discrimination, particularly under attentional cueing. The observed individual variability highlights that sensory dominance is not uniform, suggesting contributions from both bottom-up sensory reliability and top-down attentional control. Although potential semantic asymmetries between our auditory and visual stimuli may have limited perceptual binding, participants nonetheless showed multisensory enhancement. Together, these results indicate that audiovisual motion perception can approximate reliability-weighted integration but is strongly shaped by attention. Future work combining behavioral paradigms with neurophysiology will be critical for understanding the neural mechanisms supporting these dynamics and for extending models of multisensory integration to account for attention and ecological complexity.

## 4 Methods

### 4.1 Participants

Sixty healthy Vanderbilt undergraduate students and adults from the greater Nashville area (median age=19, 39 females) with self-reported normal or correct-to-normal vision, self-reported normal hearing, and no known neurological disorders participated. Undergraduate participants were compensated with course credit, and non-students were compensated monetarily at a rate of $20/hour. Each participant gave informed consent before being allowed to participate, and all recruitment procedures were in accordance with the Psychology Department Guidelines of Vanderbilt University. Basic participant demographics including sex and age were also recorded during consent. All recruitment and experimental procedures were approved by the Vanderbilt University Institutional Review Board and were carried out in accordance with the Declaration of Helsinki.

### 4.2 Stimuli

In the task, participants were presented with visual, auditory, or audiovisual stimuli. All stimuli were generated in MATLAB R2022b (The MathWorks, Inc., Natick, MA) and presented using PsychToolbox version 3 [Brainard, 1997]. Both unisensory (auditory only and visual only) and multisensory (both auditory and visual presented at the same time) stimuli were included in the experiment. Stimuli had a duration of 700 milliseconds, and were independently generated for each trial, and therefore not, “frozen.”

Visual stimuli reflected a Movshon/Newsome-type motion algorithm (Britten et al., 1992) and were displayed on a 1920×1080 cathode ray tube monitor positioned at eye-level 120 cm from the participant. The stimuli were random-dot kinematograms (RDKs) (Braddick, 1974) with 150, 3 pixel white dots (subtending <0.3° of visual space) placed randomly in a square aperture subtending a width and height of 14° visual space. The aperture size reflected the simulated displacement of the auditory stimulus at the same speed of 20°/s and a stimulus duration of 700 milliseconds. The proportion of dots that moved in a signal direction, either left or right, corresponded to the coherence value for the visual stimulus, while the remaining dots moved in random directions. Larger coherence values indicated a greater strength of motion, due to more dots within the aperture moving in the same, signal direction.

Auditory stimuli consisted of 3ms of silence followed by a 700 millisecond broad-band white noise signal (10ms rise and fall) sampled at a rate of 44100 samples per second. The signal moved at a speed of 20°/s, was embedded in partially correlated noise and played through two speakers mounted on either side of the monitor (54 cm apart) centered horizontally with the visual stimulus, on the azimuth for the participant. The auditory stimulus consisted of four different components. Independent white noise streams were presented through both speakers (100% amplitude; inter-signal correlation = 0; N1 and N2). On top of this, a combined white noise signal was presented through both speakers (100% amplitude, inter-signal correlation = 1; N3). The fourth signal stream contained the apparent motion cue, in which the sound’s amplitude faded between the two speakers from 100% to 0% over the course of one trial (N4) to create binaural motion cues. Coherence values for the auditory stimulus were the signal level, or N4 component, to noise level, N1-N3 components, ratio. Similar to the visual stimulus, larger coherence values indicated a greater strength of motion, as the N4 motion signal was louder compared to the noise. The two speakers playing the motion stimulus generated 66 dB SPL measured at the ear-level of the participant.

To estimate motion discrimination thresholds in the unisensory staircase, coherence, or the strength of the motion signal, started at 1 (100% motion signal) and was iteratively divided by 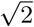, until the staircase was completed. This was to allow for a larger proportion of more difficult motion trials, where coherence was low. Coherence levels for the audiovisual conditions were identical in value and established based on psychophysical performance within a separate cohort of nine participants (four female; 19 −28 years; mean age = 23) in a pilot study. For the audiovisual cued conditions, coherence levels were 40%, 20%, 10%, and 5%. Noise trials (0% coherence or motion strength) were implemented to assess possible response biases for both unisensory and multisensory trials.

A Minolta Chroma Meter CS-100 and a sound level meter (PCB Piezotronics, Model 378C01) were used to verify the luminance and sound intensity levels, respectively. The durations of all visual and auditory stimuli were confirmed using a Hameg 507 oscilloscope (Hameg Instruments, Mainhausen, Germany) with a photovoltaic cell and microphone.

### 4.3 Paradigm

Participants were first presented with two adaptive unisensory staircase blocks (auditory-only and visual-only). The order of the unisensory staircases was randomized for each participant. Prior to beginning each block, participants were instructed to maintain fixation on a central point and report the direction of the motion stimulus (right or left) via a response box as quickly and as accurately as possible on each trial. Each unisensory trial consisted of 700 millisecond stimulus duration followed by a 1.5 – 2.5 second inter-trial interval and response window. For each unisensory block, participants began with a coherence of 100% in a random direction (left or right). A subsequent trial’s direction remained random, with an equal probability of being left or right, while the coherence depended on the previous trial’s accuracy. If correct, the next trial’s coherence had a 33% chance of being lowered by 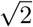 and a 66% chance of remaining the same. If incorrect, the next trial’s coherence had a 66% chance of being raised by 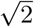 and a 33% chance of remaining the same. These unisensory trials concluded once a participant reached 30 staircase reversals, dictated by changes in coherence. Participants were given one break after 100 trials for each unisensory staircase as well as between blocks. Participants were given feedback on correctness after every trial in the form of an auditory pure tone for the two unisensory staircases. A 2000 Hz tone was played after a trial where a participant’s reported direction matched the stimulus direction (i.e., correct response), and an 800 Hz tone was played after a trial where a participant’s reported direction did not match the stimulus direction (i.e., incorrect response) or if a participant did not respond within the response time window.

After completion of the two unisensory blocks, the participants were presented with three audiovisual blocks. Each audiovisual block consisted of the same number of trials and stimuli, but the task instructions given at the beginning differed. Participants were instructed, only at the beginning of the block, to pay attention to what they heard (auditory-cued AV), what they saw (visual-cued AV), or what they saw and heard (audiovisual-cued AV) and report the direction of motion as quickly and accurately as possible. Participants completed the auditory-cued and visual-cued conditions in a random order, followed by the audiovisual-cued condition. As explained above, coherence values of 40%, 20%, 10%, and 5% were used across all audiovisual conditions for both the auditory and visual stimuli. Each audiovisual block consisted of 48 trials per coherence value, with 20 complete auditory and visual noise trials, for a total of 212 trials. Each set of 48 trials per coherence value was split into 40 congruent trials and 8 incongruent trials, evenly split between leftward and rightward trials. The auditory and visual stimuli had the same direction in congruent trials and the opposite directions in the incongruent trials, and participants were not told about the congruency of the motion. Based on a pilot study, the greater proportion of incongruent trials significantly increased the difficulty of the task. The current proportion of congruent to incongruent trials was to ease difficulty while also interrogating the incongruent trial effects. Unlike the unisensory staircases, participants were not provided with feedback for any multisensory condition. On each trial of the experiment, participants were able to respond while the stimulus was being presented or during a subsequent response period. After the 700 ms stimulus presentation, there was a response period that was randomly selected from a uniform distribution of 1.5 to 2.5 seconds (Figure 1a). The fixation cross in the middle of the CRT screen remained visible for the duration of the trial. For all trials in both the unisensory and multisensory blocks, a new trial was only initiated after the stimulus was presented and the additional response period had elapsed and was initiated whether or not the participant responded. A table showing the five conditions given to participants is shown in Figure 1b.

### 4.4 Data Analysis

Responses were collected across a range of motion coherence to assess participant performance, and these were fit to a cumulative Gaussian distribution. The slope of this distribution, where on the x-axis was leftward and rightward coherence and the y-axis was the proportion of rightward responses, was used as a measure of discrimination sensitivity for motion direction (Figure 1c). This slope was calculated as the multiplicative inverse of the standard deviation of the cumulative distribution function. Sensitivity measures were computed for each unisensory (auditory-only and visual-only) and multisensory (congruent and incongruent) condition(s) for all participants. The x-value for a 0.5 proportion rightward response was used as the point of subjective equality (PSE), a measure used to examine directional bias across participants for our task. For this measure, a negative PSE indicated a rightward bias, whereas a positive PSE indicated a leftward bias. Twelve participants were excluded from further analyses due to incomplete behavioral task data (e.g., did not complete all conditions).

Outliers in discrimination sensitivity (slope) values were assessed separately for each condition using both the median absolute deviation (MAD)–based robust z-score (> |3.5|) and the Tukey interquartile range (IQR) method (±1.5×IQR). Sensitivity values flagged by either criterion were removed from subsequent group-level analyses. Importantly, only the affected condition(s) were excluded for a given participant; remaining conditions for that participant were retained to maximize valid data contributions.

Group-level statistics first included a Shapiro-Wilk test on normality on the residuals of the distribution. Both the auditory only and incongruent auditory-cued AV group sensitivities rejected this test’s null hypothesis (*p* = 0.0102, *p <* 0.001). As such, we opted for non-parametric statistics using a repeated measures design. We first ran a Friedman test and found these group level sensitivities were significantly different (*χ*^2^ = 90.401, *p <* 0.0001). Subsequent statistical comparisons on sensitivity were Wilcoxon Signed-Rank tests with a Holm-Bonferroni correction for multiple comparisons.

### 4.5 Behavioral Modeling

Reliability-weighted models of integration were used on discrimination sensitivity to predict multisensory performance and relative sensory weights for our task. Using maximum likelihood estimation [Ernst and Banks, 2002], we computed the optimal sensory weights for each participant using the following equations:

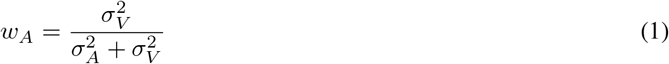

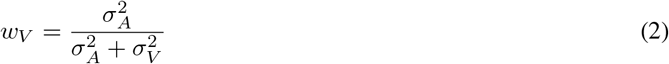

where *w*_*A*_ is the weight of the auditory cue, *w*_*V*_ is the weight of the visual cue, 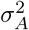 is the variance of the cumulative Gaussian distribution fit to the responses for the auditory-only task (proportional to the auditory discrimination sensitivity), and 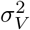 is the variance of the cumulative Gaussian distribution fit to the responses for the visual-only task (proportional to the visual discrimination sensitivity). An assumption of this model is, given these optimal weights, that the auditory and visual estimates are the only things that contribute to the multisensory estimate of motion. As such, their weights will always sum to one:

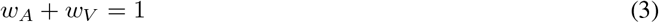

Similarly, the MLE predicted audiovisual sensitivities were derived using the responses in the unisensory conditions to predict a statistically optimal variance for the multisensory condition 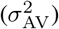:

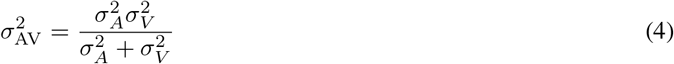

The three cued congruent AV conditions were then compared to these MLE predictions to assess the optimality of audiovisual motion integration.

Multisensory gain is commonly used to assess and quantify the degree to which two cues provide behavioral benefit beyond what is provided by the best unisensory cue [Stevenson et al., 2014]. Therefore, we calculated an enhancement index (EI) across our three cued AV conditions:

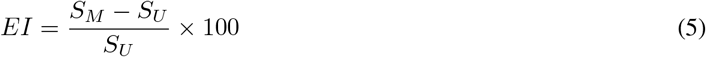

where *S*_*M*_ was the sensitivity of the auditory-cued, visual-cued, or audiovisual cued AV condition and *S*_*U*_ was the sensitivity of the best unisensory condition. This gain was used to examine relationships between multisensory gain and sensory weighting.

## 5 Author Contributions

A.J.T. contributed to study conception, study design, data collection, data analysis, figure generation, manuscript writing, and manuscript editing. K.S. contributed to data collection, data analysis, manuscript writing, and manuscript editing. R.R. and M.T.W. contributed to study conception, study design, study supervision, and manuscript editing. All authors reviewed the manuscript.

## 6 Acknowledgments

Research reported in this publication is supported by the Vanderbilt Brain Institute Trans-Institutional Programs.

The MELD consortium is supported through an unrestricted gift provided by Reality Labs Research, a division of Meta.

The authors would like to gratefully acknowledge Antonia Thelin and Adriana Schoenhaut for foundational contributions to the conception of this project. Additionally, we thank Elizabeth Brandon for her assistance with participant recruitment.

Finally, we are deeply grateful to Randolph Blake for his guidance and insight throughout this project.

## Notes

### Competing Interest Statement

The authors have declared no competing interest.

## References

D. Alais and D. Burr. No direction-specific bimodal facilitation for audiovisual motion detection. Brain Res Cogn Brain Res, 19(2):185–94, 2004a.

D. Alais and D. Burr. The ventriloquist effect results from near-optimal bimodal integration. Curr Biol, 14(3):257–62, 2004b.

A. Alink, W. Singer, and L. Muckli. Capture of auditory motion by vision is represented by an activation shift from auditory to visual motion cortex. J Neurosci, 28(11): 2690–7, 2008.

A. Alink, F. Euler, N. Kriegeskorte, W. Singer, and A. Kohler. Auditory motion direction encoding in auditory cortex and high-level visual cortex. Hum Brain Mapp, 33(4): 969–78, 2012.

R. A. Andersen. Multimodal integration for the representation of space in the posterior parietal cortex. Philos Trans R Soc Lond B Biol Sci, 352(1360):1421–8, 1997a.

R. A. Andersen. Neural mechanisms of visual motion perception in primates. Neuron, 18(6):865–72, 1997b.

D. E. Angelaki, Y. Gu, and G. C. DeAngelis. Multisensory integration: psychophysics, neurophysiology, and computation. Curr Opin Neurobiol, 19(4): 452–8, 2009.

P. W. Battaglia, R. A. Jacobs, and R. N. Aslin. Bayesian integration of visual and auditory signals for spatial localization. J Opt Soc Am A Opt Image Sci Vis, 20(7): 1391–7, 2003.

C. Battal, M. Rezk, S. Mattioni, J. Vadlamudi, and O. Collignon. Representation of auditory motion directions and sound source locations in the human planum temporale. J Neurosci, 39(12): 2208–2220, 2019.

J. K. Bizley, R. K. Maddox, and A. K. C. Lee. Defining auditory-visual objects: Behavioral tests and physiological mechanisms. Trends Neurosci, 39(2): 74–85, 2016.

S. P. Blurton, M. W. Greenlee, and M. Gondan. Cross-modal cueing in audiovisual spatial attention. Atten Percept Psychophys, 77(7): 2356–76, 2015.

David H. Brainard. The psychophysics toolbox. Spatial Vision, 10(4): 433–436, 1997.

F. Bremmer, A. Schlack, N. J. Shah, O. Zafiris, M. Kubischik, K. Hoffmann, K. Zilles, and G. R. Fink. Polymodal motion processing in posterior parietal and premotor cortex: a human fmri study strongly implies equivalencies between humans and monkeys. Neuron, 29(1): 287–96, 2001.

K. H. Britten, M. N. Shadlen, W. T. Newsome, and J. A. Movshon. The analysis of visual motion: a comparison of neuronal and psychophysical performance. J Neurosci, 12(12): 4745–65, 1992.

Laura Busse, Kenneth C. Roberts, Roy E. Crist, Daniel H. Weissman, and Marty G. Woldorff. The spread of attention across modalities and space in a multisensory object. Proceedings of the National Academy of Sciences, 102(51): 18751–18756, 2005.

C. Cappe, G. Thut, V. Romei, and M. M. Murray. Selective integration of auditory-visual looming cues by humans. Neuropsychologia, 47(4): 1045–52, 2009.

C. Cappe, A. Thelen, V. Romei, G. Thut, and M. M. Murray. Looming signals reveal synergistic principles of multisensory integration. J Neurosci, 32(4): 1171–82, 2012.

S. Carlile and J. Leung. The perception of auditory motion. Trends Hear, 20, 2016.

T. A. Chaplin, B. J. Allitt, M. A. Hagan, M. G. P. Rosa, R. Rajan, and L. L. Lui. Auditory motion does not modulate spiking activity in the middle temporal and medial superior temporal visual areas. Eur J Neurosci, 48(4):2013–2029, 2018a.

T. A. Chaplin, M. G. P. Rosa, and L. L. Lui. Auditory and visual motion processing and integration in the primate cerebral cortex. Front Neural Circuits, 12:93, 2018b.

Sarah E. Donohue, Kenneth C. Roberts, Tineke Grent-’t Jong, and Marty G. Woldorff. The cross-modal spread of attention reveals differential constraints for the temporal and spatial linking of visual and auditory stimulus events. The Journal of Neuroscience, 31(22): 7982–7990, 2011.

A. Duyar, A. Pavan, and H. Kafaligonul. Attentional modulations of audiovisual interactions in apparent motion: Temporal ventriloquism effects on perceived visual speed. Atten Percept Psychophys, 84(7): 2167–2185, 2022.

M. O. Ernst and M. S. Banks. Humans integrate visual and haptic information in a statistically optimal fashion. Nature, 415(6870): 429–33, 2002.

M. O. Ernst and H. H. Bulthoff. Merging the senses into a robust percept. Trends Cogn Sci, 8(4): 162–9, 2004.

C. R. Fetsch, A. Pouget, G. C. DeAngelis, and D. E. Angelaki. Neural correlates of reliability-based cue weighting during multisensory integration. Nat Neurosci, 15(1): 146–54, 2011.

T. C. Freeman, J. Leung, E. Wufong, E. Orchard-Mills, S. Carlile, and D. Alais. Discrimination contours for moving sounds reveal duration and distance cues dominate auditory speed perception. PLoS One, 9(7):e102864, 2014.

M. C. Hagen, O. Franzen, F. McGlone, G. Essick, C. Dancer, and J. V. Pardo. Tactile motion activates the human middle temporal/v5 (mt/v5) complex. Eur J Neurosci, 16(5): 957–64, 2002.

J. Hocking and C. J. Price. The role of the posterior superior temporal sulcus in audiovisual processing. Cereb Cortex, 18(10): 2439–49, 2008.

Y. Kamitani and F. Tong. Decoding seen and attended motion directions from activity in the human visual cortex. Curr Biol, 16(11): 1096–102, 2006.

C. Kayser, C. I. Petkov, M. Augath, and N. K. Logothetis. Functional imaging reveals visual modulation of specific fields in auditory cortex. J Neurosci, 27(8): 1824–35, 2007.

S. J. Kayser and C. Kayser. Shared physiological correlates of multisensory and expectation-based facilitation. eNeuro, 7(2), 2020.

S. J. Kayser, M. G. Philiastides, and C. Kayser. Sounds facilitate visual motion discrimination via the enhancement of late occipital visual representations. Neuroimage, 148: 31–41, 2017.

K. Krumbholz, M. Schonwiesner, R. Rubsamen, K. Zilles, G. R. Fink, and D. Y. von Cramon. Hierarchical processing of sound location and motion in the human brainstem and planum temporale. Eur J Neurosci, 21(1): 230–8, 2005.

R. Lewis and U. Noppeney. Audiovisual synchrony improves motion discrimination via enhanced connectivity between early visual and auditory areas. J Neurosci, 30(37): 12329–39, 2010.

S. Majerus, F. Peters, M. Bouffier, N. Cowan, and C. Phillips. The dorsal attention network reflects both encoding load and top-down control during working memory. J Cogn Neurosci, 30(2): 144–159, 2018.

J. H. Maunsell and D. C. Van Essen. Functional properties of neurons in middle temporal visual area of the macaque monkey. i. selectivity for stimulus direction, speed, and orientation. J Neurophysiol, 49(5): 1127–47, 1983.

M. E. McCourt and L. M. Leone. Auditory capture of visual motion: effects on perception and discrimination. Neuroreport, 27(14): 1095–100, 2016.

G. F. Meyer and S. M. Wuerger. Cross-modal integration of auditory and visual motion signals. Neuroreport, 12(11): 2557–60, 2001.

S. Molholm, P. Sehatpour, A. D. Mehta, M. Shpaner, M. Gomez-Ramirez, S. Ortigue, J. P. Dyke, T. H. Schwartz, and J. J. Foxe. Audio-visual multisensory integration in superior parietal lobule revealed by human intracranial recordings. J Neurophysiol, 96(2): 721–9, 2006.

Michael L. Morgan, Gregory C. DeAngelis, and Dora E. Angelaki. Multisensory integration in macaque visual cortex depends on cue reliability. Neuron, 59(4): 662–673, 2008.

C. Poirier, O. Collignon, A. G. Devolder, L. Renier, A. Vanlierde, D. Tranduy, and C. Scheiber. Specific activation of the v5 brain area by auditory motion processing: an fmri study. Brain Res Cogn Brain Res, 25(3): 650–8, 2005.

R. Ptak and A. Schnider. The dorsal attention network mediates orienting toward behaviorally relevant stimuli in spatial neglect. J Neurosci, 30(38): 12557–65, 2010.

A. Rajan, S. Meyyappan, Y. Liu, I. B. H. Samuel, B. Nandi, G. R. Mangun, and M. Ding. The microstructure of attentional control in the dorsal attention network. J Cogn Neurosci, 33(6): 965–983, 2021.

M. Rezk, S. Cattoir, C. Battal, V. Occelli, S. Mattioni, and O. Collignon. Shared representation of visual and auditory motion directions in the human middle-temporal cortex. Curr Biol, 30(12):2289–2299 e8, 2020.

T. Rohe and U. Noppeney. Reliability-weighted integration of audiovisual signals can be modulated by top-down attention. eNeuro, 5(1), 2018.

R. J. Snowden and T. C. Freeman. The visual perception of motion. Curr Biol, 14(19):R828–31, 2004.

S. Soto-Faraco, A. Kingstone, and C. Spence. Multisensory contributions to the perception of motion. Neuropsychologia, 41(13): 1847–62, 2003.

S. Soto-Faraco, C. Spence, and A. Kingstone. Cross-modal dynamic capture: congruency effects in the perception of motion across sensory modalities. J Exp Psychol Hum Percept Perform, 30(2): 330–45, 2004.

R. A. Stevenson and T. W. James. Audiovisual integration in human superior temporal sulcus: Inverse effectiveness and the neural processing of speech and object recognition. Neuroimage, 44(3): 1210–23, 2009.

R. A. Stevenson, D. Ghose, J. K. Fister, D. K. Sarko, N. A. Altieri, A. R. Nidiffer, L. R. Kurela, J. K. Siemann, T. W. James, and M. T. Wallace. Identifying and quantifying multisensory integration: a tutorial review. Brain Topogr, 27 (6):707–30, 2014.

D. Talsma, D. Senkowski, S. Soto-Faraco, and M. G. Woldorff. The multifaceted interplay between attention and multisensory integration. Trends Cogn Sci, 14(9): 400–10, 2010.

M. T. Wallace, G. E. Roberson, W. D. Hairston, B. E. Stein, J. W. Vaughan, and J. A. Schirillo. Unifying multisensory signals across time and space. Exp Brain Res, 158(2): 252–8, 2004.

Robert B. Welch and David H. Warren. Immediate perceptual response to intersensory discrepancy. Psychological Bulletin, 88(3): 638–667, 1980.

S. Werner and U. Noppeney. Distinct functional contributions of primary sensory and association areas to audiovisual integration in object categorization. J Neurosci, 30(7): 2662–75, 2010.

